# Assessment of the macrovascular contribution to resting-state fMRI functional connectivity at 3 Tesla

**DOI:** 10.1101/2023.10.19.563131

**Authors:** Xiaole Z. Zhong, Yunjie Tong, J. Jean Chen

## Abstract

In resting-state functional magnetic resonance imaging (rs-fMRI) functional connectivity (FC) mapping, temporal correlation are widely assumed to reflect synchronized neural-related activaty. Although a large number of studies have demonstrated the potential vascular effects on FC, little research has been conducted on FC resulting from macrovascular signal fluctuations. Previously, Tong et al. found (Tong et al., 2019b) a robust anti-correlation between the fMRI signals in the internal carotid artery and the internal jugular vein (and the sagittal sinus). The present study extends the previous study to include all detectable major veins and arteries in the brain in a systematic analysis of the macrovascular contribution to the functional connectivity of the whole-GM. This study demonstrates that: 1) The macrovasculature consistently exhibited strong correlational connectivity among itself, with the sign of the correlations varying between arterial and venous connectivity; 2) GM connectivity was found to have a strong macrovascular contribution, stronger from veins than arteries; 3) functional connectivity originating from the macrovasculature displayed disproportionately high spatial variability compared to across all GM voxels; 4) macrovascular contributions to connectivity were still evident well beyond the confines of the macrovascular space. These findings highlight the extensive contribution to rs-fMRI BOLD and FC predominantly by large veins, but also by large arteries. These findings pave the way for future studies aimed at more comprehensively modeling and thereby removing these macrovascular contributions.

## Introduction

Resting-state blood-oxygenation level-dependent (BOLD) functional magnetic resonance imaging (rs-fMRI) is extensively used for mapping resting-state functional connectivity (rs-fcMRI) and for providing brain-health assessments (Srivastava et al., 2022; van den Heuvel and Hulshoff Pol, 2010). This technique has been used in aging, Alzheimer’s disease (Vemuri et al., 2012) and stroke (Ovadia-Caro et al., 2014), among others, and is commonly applied in studying brain development and aging processes (Andrews-Hanna et al., 2007; Geerligs et al., 2015). Most commonly, rs-fcMRI is calculated by correlation analysis (Bandettini et al., 1993; Biswal et al., 1995), in which the BOLD signal from seed voxels are correlated with those in the rest of the brain. Correlation is commonly interpreted as synchronous neural activity. It is possible, however, for large blood vessels in the brain to exhibit strong correlations among themselves as well as the global mean BOLD signal (Tong et al., 2019b).As the macrovascular system is responsible for supplying and draining blood from a large area of the brain, it is unlikely to be associated with local specific neural activity (Uludağ and Blinder, 2018), but rather with physiological noise and systemic processes. Thus, functional connectivity (FC) estimates may be biased by these macrovascular correlations (Huck et al., 2023; Kalcher et al., 2015). In order to understand macrovascular bias on FC estimates using rs-fMRI, it is imperative to understand the nature of signals that they carry, and the magntitude of their resultant BOLD signals.

The debate between vein versus brain in task-based fMRI has a long history. At a very early age, with both rodent experiments and simulation studies, Boxerman and colleagues demonstrated that macrovasculature affects gradient-echo (GRE) BOLD contrast more than microvasculature (Boxerman et al., 1995). Although it is true that the sensitivity of microvasculature increases with an increase in the main magnetic field (Menon and Goodyear, 1999), veins still account for the majority of the signal fluctuation in BOLD signals at 7T (Menon, 2012). In previous studies with rats, it was demonstrated that fcMRI could detect venous structures (Hyde and Li, 2014) and that 11.7 T MRI could detect veins twice as well as brain tissue (Yu et al., 2012). Several studies focusing on the penetrating arteries and ascending veins have suggested, although they are much smaller in size than the macrovasculature used in this study, that it is not correlated with local neural activity (Uludağ and Blinder, 2018) and that multiple factors (for example, orientation and cortical depth of the blood vessel) are involved (Viessmann et al., 2019) that make it difficult to comprehend. It is also possible that other factors, such as spatial resolution and participant positioning, may have an effect on the macrovascular effect. It is unclear how these factors interact with one another.

Unlike in conventional task-based fMRI, it is common in rs-fMRI to focus almost entirely on the dynamic BOLD signal in the low-frequency band, due to established knowledge regarding the frequency of the canonical hemodynamic response (Tong et al., 2019a). Correspondingly, most rs-fcMRI studies have focused on such low-frequency BOLD fluctuations (Biswal et al., 1995; Josephs and Henson, 1999). Within this frequency range, a large body of work has demonstrated consistent time-lagged spatial correlational structures stemming from what has been termed systemic low-frequency oscillations (sLFOs) (Tong et al., 2015). These sLFOs permeate the brain volume, being particularly pronounced in the vasculature. The most commonly referenced contributors to the sLFOs would be vasomotion (Hundley et al., 1988; Mayhew et al., 1996; Rivadulla et al., 2011), heart rate (Thayer et al., 2012), respiratory volume variability (Birn et al., 2006; Chang et al., 2009), gastric oscillations (Mohamed Yacin et al., 2011; Rebollo et al., 2018) and variations in carbon dioxide levels (Sassaroli et al., 2012; Wise et al., 2004). Notably, previous observations of robust anti-correlations between arterial and venous BOLD signals in the low-frequency range (Tong et al., 2019b) demonstrated the contribution of arterial BOLD to the rs-fMRI correlational structure, overturning the long-held belief that arterial BOLD effects are negligible. Despite the diminutive arterial contrast, its contribution to rs-fMRI is highlighted by the fact that correlation-based rs-fcMRI metrics can be strongly influenced by low-amplitude and yet highly-synchronous BOLD signal variations.

Given that the macrovascular BOLD signal is deemed not neuronal specific, some may argue that it would be simple to identify and mask out regions with macrovasculature during rs-fcMRI analysis. However, according to the biophysical model of BOLD signal, the T2* relaxation effect caused by the susceptibility differences between blood and brain tissue originating from macrovasculature is capable of extending well beyond the voxel (Ogawa et al., 1993b), as recently shown in an in-vivo experiment (Huck et al., 2023). Moreover, previous results indicated that macrovascular signals and global BOLD signals are highly synchronous, given by their high cross-correlation coefficients (Tong et al., 2015), suggesting that macrovasculature may likely contribute significantly to the gray matter (GM) BOLD signal beyond the intravascular space. The potential spatial extent of macrovascular contribution to rs-fcMRI metrics in the perivascular space elevates the potential influence of macrovascular BOLD on rs-fcMRI and the challenge in removing it, and yet, it is not well understood..

Indeed, this study was inspired by previous observations of robust correlations between arterial and venous signals in the low-frequency range (Tong et al., 2015). More recently, Huck et al. (Huck et al., 2023) extended the investigation into the rest of the venous vasculature and found that macrovascular contributions were significant for rs-fcMRI metrics. Although these two previous studies are informative, they either only examine the correlation between a very limited number of large blood vessels rather than the entire vasculature (Tong et al., 2015) or only examine venous vasculature without examining arterial vasculature (Huck et al., 2023). In addition, the macrovascular effect present in the perivascular space, despite its importance, has largely been overlooked. In spite of the fact that Huck et al included perivascular space in their analysis, their results were heavily dependent on fitting high-order parameters, which may not fully capture the macrovascular contribution to perivascular space. The purpose of this study was to demonstrate macrovascular effects in both the macrovasculature and the perivascular space by utilizing time-of-flight imaging (TOF) at 3 Tesla to locate macrovasculature.

## Methods

### Participants

This study used data from the Midnight Scan Club (MSC) dataset, which comprises of MRI data from ten young, healthy, right-handed participants, five males and five females, ages 24-34 (Gordon et al., 2017). The study protocol was approved by the Human Studies Committee and Institute Review Board at Washington University School of Medicine in accordance with the Declaration of Helsinki. This data can be obtained from the OpenNeuro database, with accession number ds000224.

### MRI acquisition

Subjects underwent twelve imaging sessions on a Siemens TRIO 3T MRI scanner (Siemens Healthcare GmbH, Erlangen, Germany) on separate days. In this section, only the protocols related to this study are listed.

In total, four T1-weighted data sets (sagittal, 224 slices, 0.8 mm isotropic resolution, TE = 3.74 ms, TR = 2400 ms, TI = 1000 ms, flip angle = 8°), four angiograms (transverse, 0.6 x 0.6 x 1.0 mm, 44 slices, TR = 25ms, TE = 3.34ms), eight venograms (four sagittal scans: 0.8 x 0.8 x 2.0 mm thickness, 120 slices, TR = 27 ms, TE = 7.05 ms; four coronal scans: 0.7 x 0.7 x 2.5 mm thickness, 128 slices, TR = 28 ms, TE = 7.18 ms), and 10 sessions of rs-fMRI with a gradient-echo EPI sequence (TR = 2.2 s, TE = 27 ms, flip angle = 90 degrees, 4 mm isotropic resolution, 36 slices, scan time = 30 minutes) were collected for each participant. The participants were instructed to fixate on a white crosshair on a black background during the rs-fMRI scans. An EyeLink 1000 eye-tracking system (SR-Research, Ottawa, Canada, http://www.sr-research.com) was used to ensure participant alertness during the scans.

### Blood vessel segmentation and data processing

The strategies for macrovasculature segmentation and processing are summarized in **Figure 1a**. Angiograms and venograms were registered to T1 space (FSL MCFLIRT) and segmented using the Brain Charter Toolbox (Bernier et al., 2018). Visual inspection was performed to ensure the absence of artifacts. Vascular segmentations from all angiograms and venograms across all encoding directions were summed and then binarized to produce the final vascular segmentation. This was in turn downsampled to the rs-fMRI resolution. Due to the fact that both angiograms and venograms contained both arteries and veins, each arterial and venous map was manually separated after downsampling into different maps.

**Figure 1.**
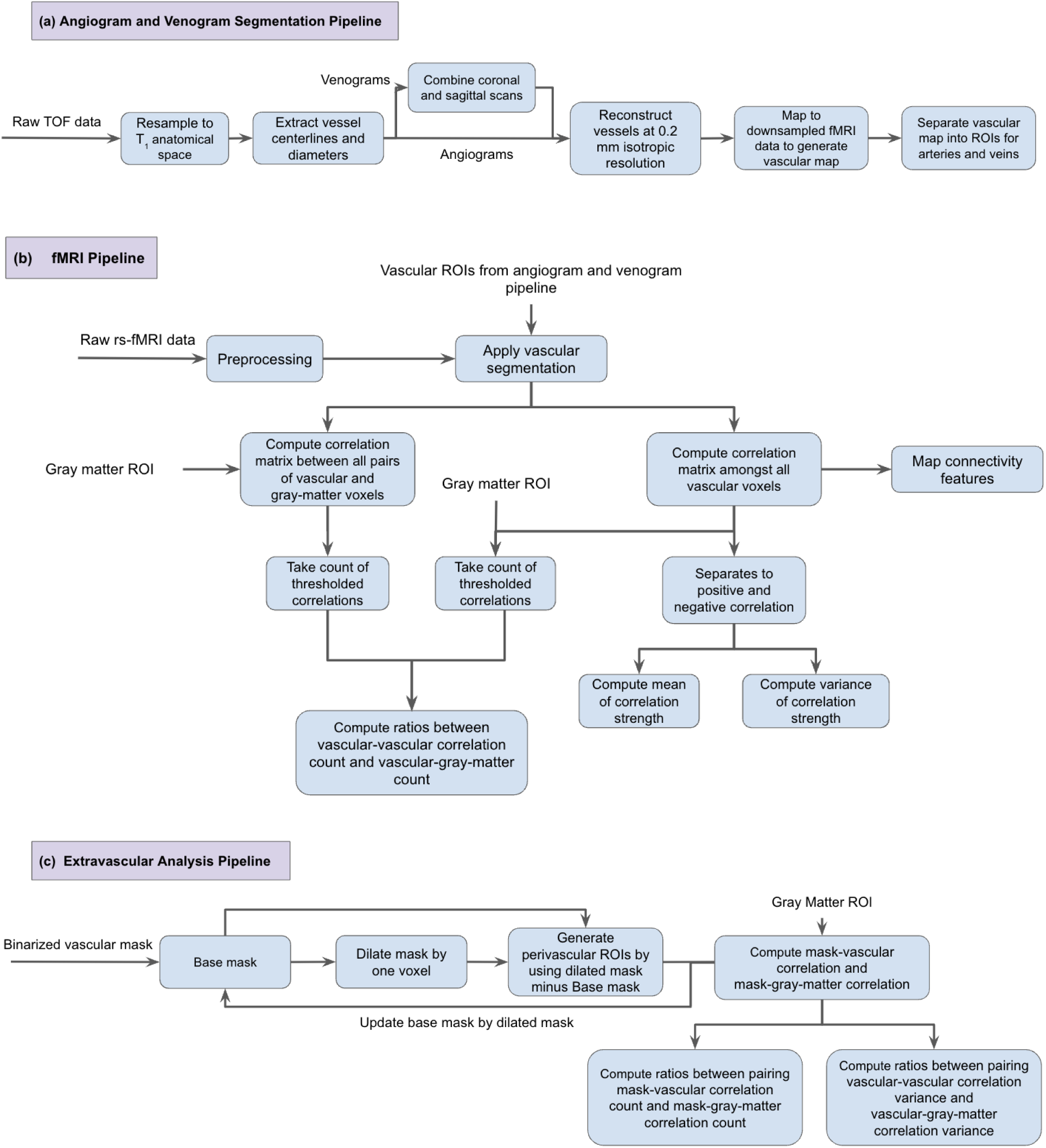
Overview of the analysis procedure. a) the angiogram and venogram preprocessing and segmentation pipeline; b) the fMRI analysis pipeline; c) the analysis pipeline for assessing the extravascular effects.

**Figure 2.**
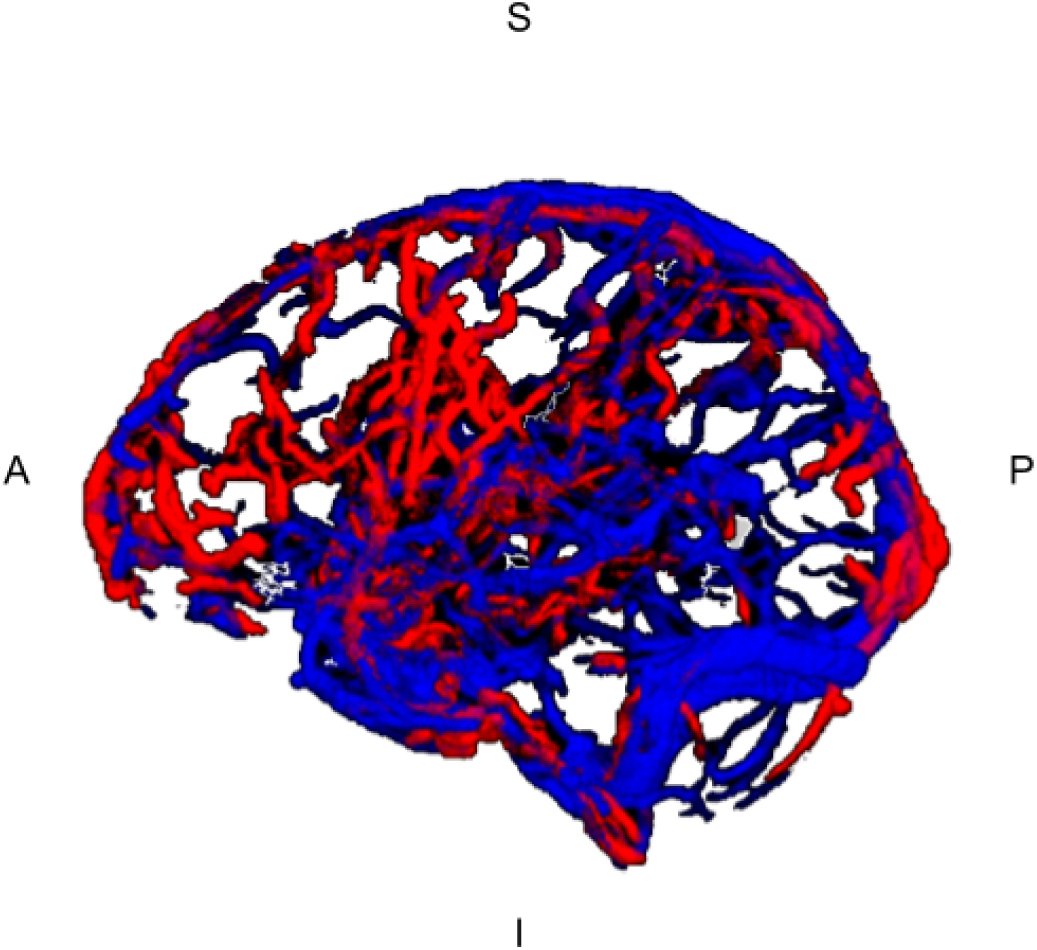
Sample segmented blood vessels (Before separating the venous and arterial manually). Red: segmented based on MRAs; blue: segmented based on MRVs. I = inferior, S = superior, A = anterior, P = posterior.

### fMRI processing and FC analysis

A summary of the rs-fMRI processing procedures can be found in **Figure 1b**. fMRI preprocessing pipeline was implemented with tools from FSL (Jenkinson et al., 2012), AFNI (Cox, 1996) and FreeSurfer (Fischl, 2012). The following steps were included in the preprocessing steps: (a) 3D motion correction (FSL MCFLIRT), (b) slice-timing correction (FSL slicetimer), (c) brain extraction (FSL bet2 and FreeSurfer mri_watershed), (d) rigid body coregistration of functional data to the individual T1 image (FSL FLIRT), (e) regression of the mean signals from white-matter (WM) and cerebrospinal fluid (CSF) regions (fsl_glm), (f) bandpass filtering to obtain frequency band 0.01-0.1 Hz (AFNI 3dBandpass), and (g) the data were spatially smoothed with 6 mm full-width half-maximum (FWHM) Gaussian kernel (FSL fslmaths).

For each session of each participant, the voxel-wise vascular-driven connectivity metrics were calculated for each pair of voxels, namely (1) venous-venous correlation, defined as venous voxel correlated with all other venous voxels; (2) arterial-arterial correlation, defined as arterial voxel correlated with all other arterial voxels; and (3) arterial-venous correlation, defined as arterial voxel correlated with all venous voxels (and transposed to obtained venous-arterial correlation). Then, inspired by previous research (Buckner et al., 2009; Cole et al., 2010), we also computed the degree of connectivity (D) and strength of connectivity (S) as global metrics of connectivity. At each vascular voxel, D was computed by counting the number of voxels connected to each other vascular voxel and having a correlation coefficient greater than 0.15 (Buckner et al., 2009; Cole et al., 2010). Voxelwise, connectivity strength (S) and spatial variance (σ^2^) are computed as the mean and variance of these thresholded correlations. The maps of D, S and σ^2^ maps were further separated into:

- D_V,V_, S_V,V_, σ^2^_V,V_: based solely on venous-venous correlations;
- D_V,A_, S_V,A_, σ^2^_V,A_: based solely on venous-arterial correlations (venous-arterial correlation, computed within the venous vasculature);
- D_A,V,_ S_A,V_, σ^2^_A,V_: based solely on arterial-venous correlations (arterial-venous correlation, computed within the arterial vasculature);
- D_A,A_, S_A,A_, σ^2^_A,A_: based solely on arterial-arterial correlation.

Moreover, correlation-strength maps are separated by positive and negative correlations.

### Macrovascular contribution to rs-fcMRI

For a better understanding of the macrovascular contribution to overall brain connectivity, we also computed GM degree of connectivity, defined as the number of correlations between each GM voxel and each other GM voxel. These correlations were thresholded at 0.15, as mentioned earlier. GM masks were applied on top of the macrovascular mask and maped consisting of the ratios of macrovascular to whole-GM degrees of connectivity. Additionally, variance ratios were calculated by substituting variance of connectivity for degree of connectivity. These ratios were further separated into:

- Arterial-arterial, in terms of D_A,A_:D_A,GM_, σ^2^_A,A_: σ^2^_A,GM_;
- Arterial vs. venous, in terms of D_A,V_:D_A,GM_, σ^2^_A,V_: σ^2^_A,GM_D_V,A_:D_V,GM_, σ^2^_V,A_: σ^2^_A,GM_;
- Venous-venous, in terms of D_V,V_:D_V,GM_, σ^2^_V,V_: σ^2^_V,GM_.

### Perivascular effects on rs-fcMRI metrics

Figure 1c illustrates the methods used to assess the perivascular contributions by the macrovasculature. Briefly, we expanded the analysis from the previous section into the extravascular space. That is, we computed voxel-wise pairwise correlations stemming from perivascular voxels for arterial and venous macrovascular voxels (similar to the case of arterial and venous correlations described earlier), and then derived degrees of correlation from these perivascular voxels. The perivascular ROIs were generated as follows. First, the arterial and venous segmentations were dilated by 1 voxel in 3D. Second, the original vascular ROI was subtracted from the dilated vascular ROI. This produced the extravascular ROI that is 1 voxel distant from the vessel. This process was repeated for perivascular distances of 2 and 3 voxels. With the same approach as that used for the case of arterial and venous correlations, we computed ratios of macrovascular to whole-GM degrees (D_A(ex),A_:D_A(ex),GM_ for the ratio between arterial perivascular space to arterial correlation degree and arterial perivascular space to GM correlation degree and D_V(ex),V_:D_V(ex),GM_ for the ratio between venous perivascular space to venous correlation degree and venous perivascular space to GM correlation degree). Additionally, computed ratios of macrovascular to whole-GM variances (σ^2^_A(ex),A_:σ^2^_A(ex),GM_, for the ratio between arterial perivascular space to arterial correlation variance and arterial perivascular space to GM correlation variance and σ^2^:σ^2^ for the ratio between venous perivascular space to venous correlation variance and venous perivascular space to GM correlation variance) by substituting σ^2^ for the D in the computation of degree ratio maps mentioned previously.

## Results

### Macrovascular connectivity

In order to assess the connectivity arising from macrovasculature, three metrics were used: degree of connectivity (D) (Fig. 3), strength of connectivity (S) (Fig. 4) and variance of connectivity (σ^2^) (Fig. 5).

**Figure 3.**
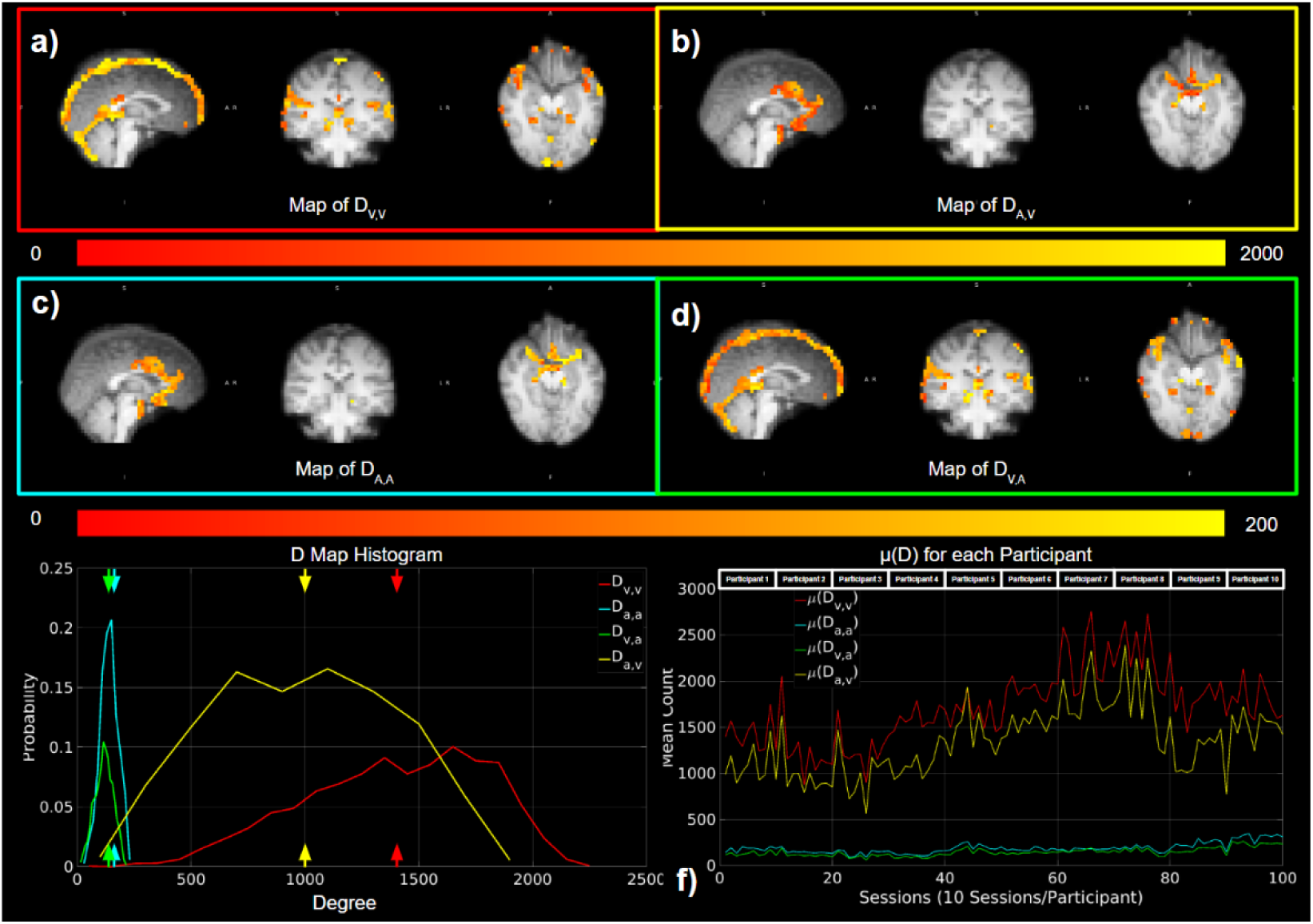
Degrees of macrovascular connectivity. Red indicates D_V,V_; yellow indicates D_A,V_; cyan indicates D_A,A_; green indicates D_V,A_. (a)-(e) represent results from a representative data set, while (f) shows results from all data sets.

**Figure 4.**
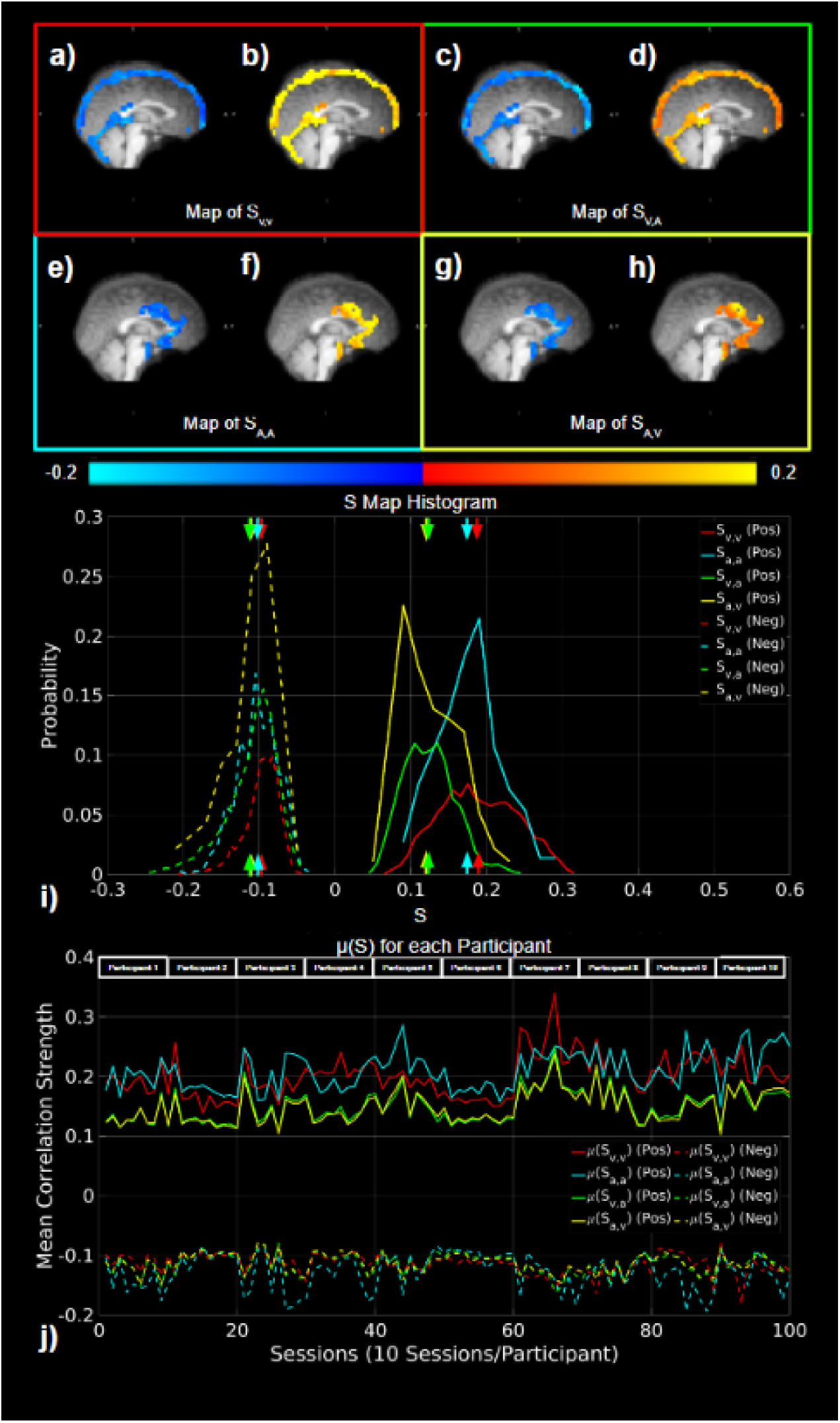
Strengths of macrovascular connectivity. Red indicates S_V,V_; yellow indicates S_A,V_; cyan indicates S_A,A_; green indicates S_V,A_. (a)-(i) represent results from a representative data set, while (j) shows results from all data sets.

**Figure 5.**
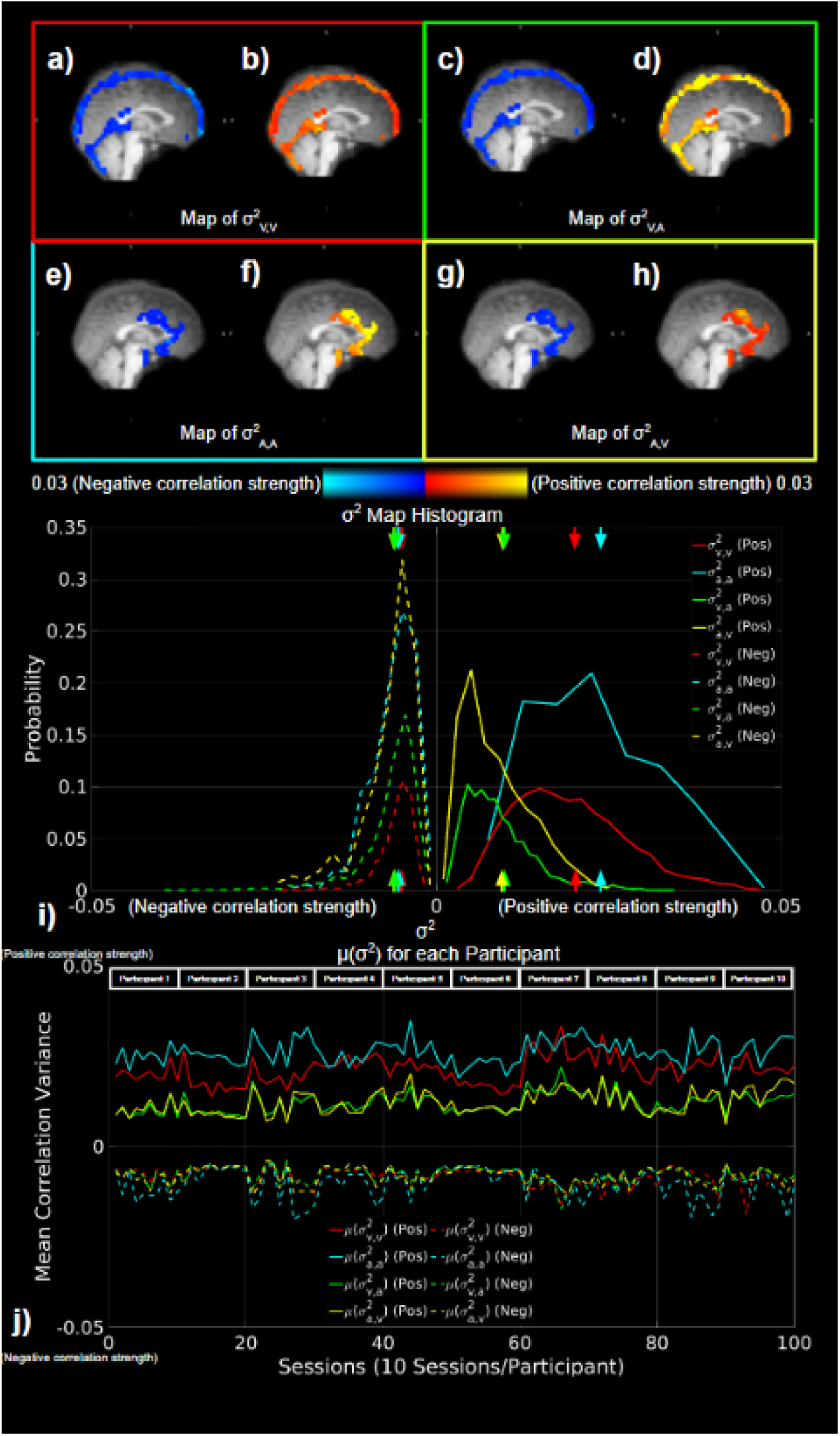
Variance of macrovascular connectivity. Red indicates σ^2^_v,v_; yellow indicates σ^2^_A,V_; cyan indicates σ^2^_A,A_; green indicates σ^2^_V,A_, with correlation with positive S represented by solid lines, and correlation with negative S represented by dashed lines. (a)-(i) represent results from a representative data set, while (j) shows results from all data sets.

### Degree of macrovascular connectivity

In Figure 3, we demonstrated strong apparent FC within the macrovascular regions based on degree of connectivity. As seen in the results from a representative dataset, voxels with non-zero degrees of vascular-related connections (D_V,V,_ D_A,A_, D_A,V_ and D_V,A_) were found to span the entire macrovascular ROI, with higher values at the site of larger vessels (e.g. the superior sagittal sinus and Circle of Willis) (Fig. 3a**-d**). The histograms of D values show that D_V,V_ and D_A,V_ are higher than D_V,A_ and D_A,A_, with the maximum degree exceeding 2000 (Fig. 3e). These patterns are also evident in μ(D) for these four correlation categories across all participants in all scan sessions (Fig. 3f).

There was also evidence of strong apparent connectivity within macrovasculature based on the strengths and variances metrics (Fig. 4 **and** Fig. 5). As seen in these results from the same representative dataset as used in the previous figures, higher S_V,V_ magnitudes are associated with positive than negative correlations (Fig. 4a**,b**). Likewise, higher S_A,A_ magnitudes are also associated with positive than negative correlations (Fig. 4e**,f**). Conversely, S_V,A_ was similar in magnitude for both negative and positive correlations (Fig. 4c, d, g, h). These differences are further illustrated by the whole-vasculature histograms with S_A,A_ and S_V,V_ show higher magnitudes than S_A,V_ and S_V,A_ (Fig. 4i). Moreover, as shown across all subjects (Fig. 4j), the strong S_A,A_ and S_V,V_ patterns are observed across all participants in all scan sessions, outweighting values for S_A,V_ and S_V,A_.

Similar to the mean correlation strength S, the correlation variance σ^2^ and σ^2^ are higher for positive than negative correlations, as shown for the same representative data set as in previous figures (Fig. 5a**,b****,e,f**). There was no apparent difference in σ^2^ and σ^2^ between positive and negative correlations (Fig. 5c**,d****,g,h**). Moreover, the histograms with distribution probability showed the same patterns as maps, which suggest (Fig. 5i), a pattern that is consistent across all sessions from all participants exhibited the same pattern (Fig. 5j).

### Contribution of the macrovascular connectivity to GM functional connectivity

The contribution of macrovascular to whole-GM FC was first quantified as ratios between vascular and GM degrees of connectivity (D_V,V_:D_V,GM_, D_V,A_:D_v,GM_, D_A,V_:D_A,GM_, and D_A,A_:D_A,GM_) (Fig. 6). According to the values we obtained for venous contributions through D_v,v_:D_v,GM_ (Fig. 6a) and D_A,V_:D_A,GM_ (Fig. 6d), large portion of the significant correlation in D_GM_ could come from the vascular ROIs. This fraction could reach up to 48.8% and 45.1%, respectively. The arterial ratios D_A,A_:D_A,GM_ (Fig. 6c) and D_V,A_:D_V,GM_ (Fig. 6b) are lower, but still as high as 7.98% and 16.2%, respectively. These differences are further illustrated by the histograms, which show the degree ratios to be highest for D_V,V_:D_V,GM_, followed by D_A,V_:D_A,GM_, D_A,A_:D_A,GM_, and the lowest for D_V,A_:D_V,GM_ (Fig. 6e). These patterns were also evident in mean degree ratios for the above four groups of correlations across all participants in all scan sessions (Fig. 6f). Furthermore, across all sessions for all participants, the mean degree ratios μ(D_V,V_:D_V,GM_) and μ(D_A,A_:D_A,GM_) fluctuated between 12% and 29%, while the μ(D_A,A_:D_V,GM_) and μ(D_V,A_:D_A,GM_) were less than 5%.

**Figure 6.**
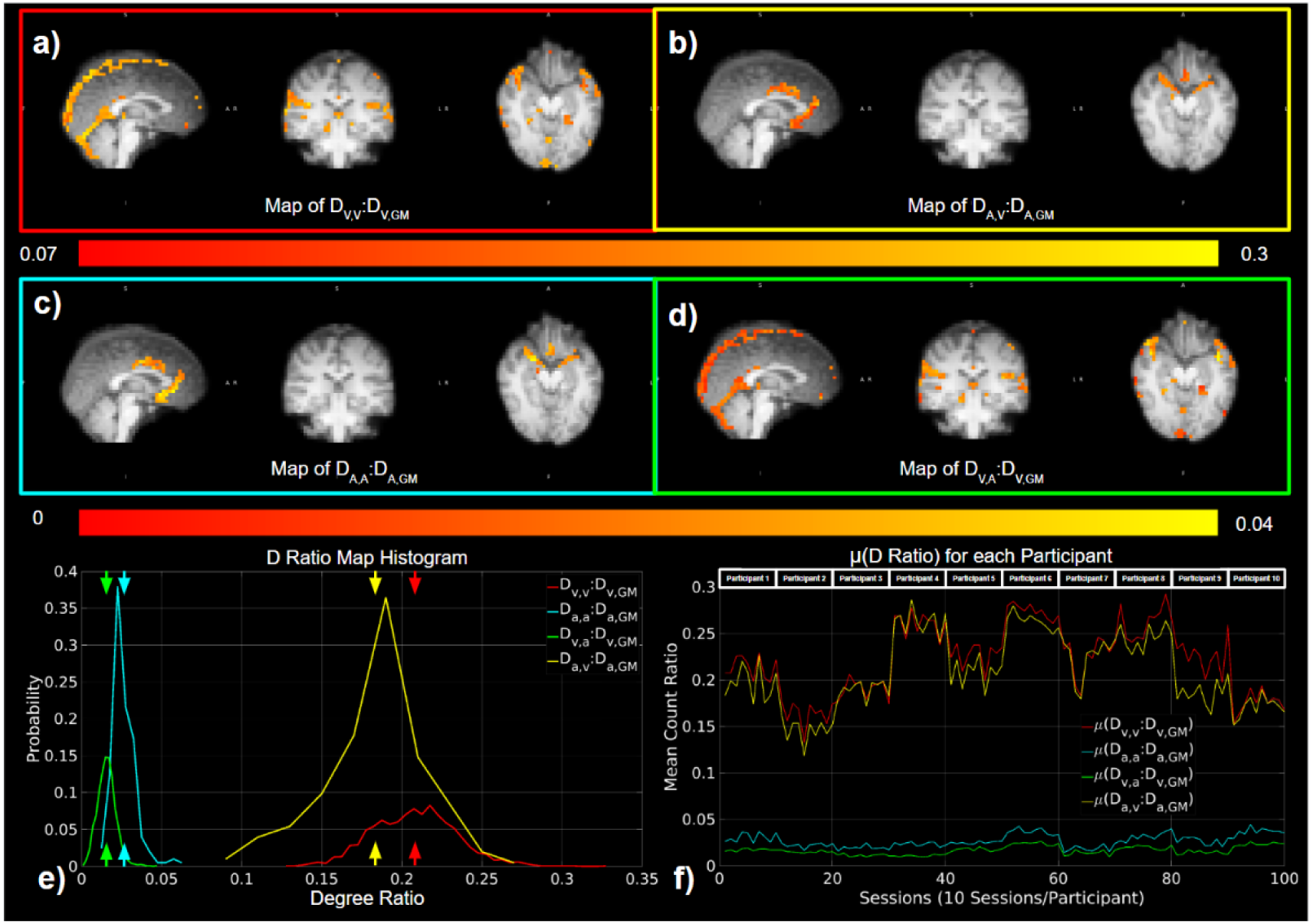
Contribution of the macrovascular connectivity to overall tissue degree of connectivity. The contribution was quantified as ratios between vascular and tissue degrees of connectivity, e.g. D_v,v_:D_v,GM_, D_v,a_:D_v,GM_, D_a,v_:D_a,GM_, and D_a,a_:D_a,GM_. (a)-(e) represent results from a representative data set, while (f) shows results from all data sets. Red indicates D_v,v_:D_v,GM_; yellow indicates D_a,v_:D_a,GM_; cyan indicates D_a,a_:D_a,GM_; green indicates D_v,a_:D_v,GM_.

As a measure of the spatial variability of functional connectivity associated with the macrovasculature relative to that of whole GM, variance ratios (σ^2^_V,V_:σ^2^_V,GM_, σ^2^_V,A_:σ^2^_v,GM_, σ^2^_A,V_:σ^2^_A,GM_, and σ^2^_A,A_:σ^2^_A,GM_) between vascular and GM degrees of connectivity were computed. We found that the variance ratio σ^2^_A,A_:σ^2^_A,GM_ (Fig. 7a) exhibited the highest values, reaching up to 3.73. Followed by σ^2^_V,A_:σ^2^_v,GM_ which reached up to ∼3.05 (Fig. 7b). σ^2^_V,V_:σ^2^_v,GM_ (Fig. 7c) and σ^2^_A,V_:σ^2^_A,GM_ (Fig. 7d) exhibited lower values but still could be as high as 2.07 and 1.45, respectively. According to the histogram (Fig. 7e), the μ(σ^2^_V,V_:σ^2^_v,GM_) and μ(σ^2^_V,V_:σ^2^_v,GM_) had slightly different maxima, with μ(σ^2^_V,V_:σ^2^_v,GM_) higher than the μ(σ^2^_V,V_:σ^2^_v,GM_). Across all sessions for all participants, we observed that the μ(σ^2^_A,A_:σ^2^_A,GM_) was consistently higher than variance ratios for any connectivity involved veins (Fig. 7f).

**Figure 7.**
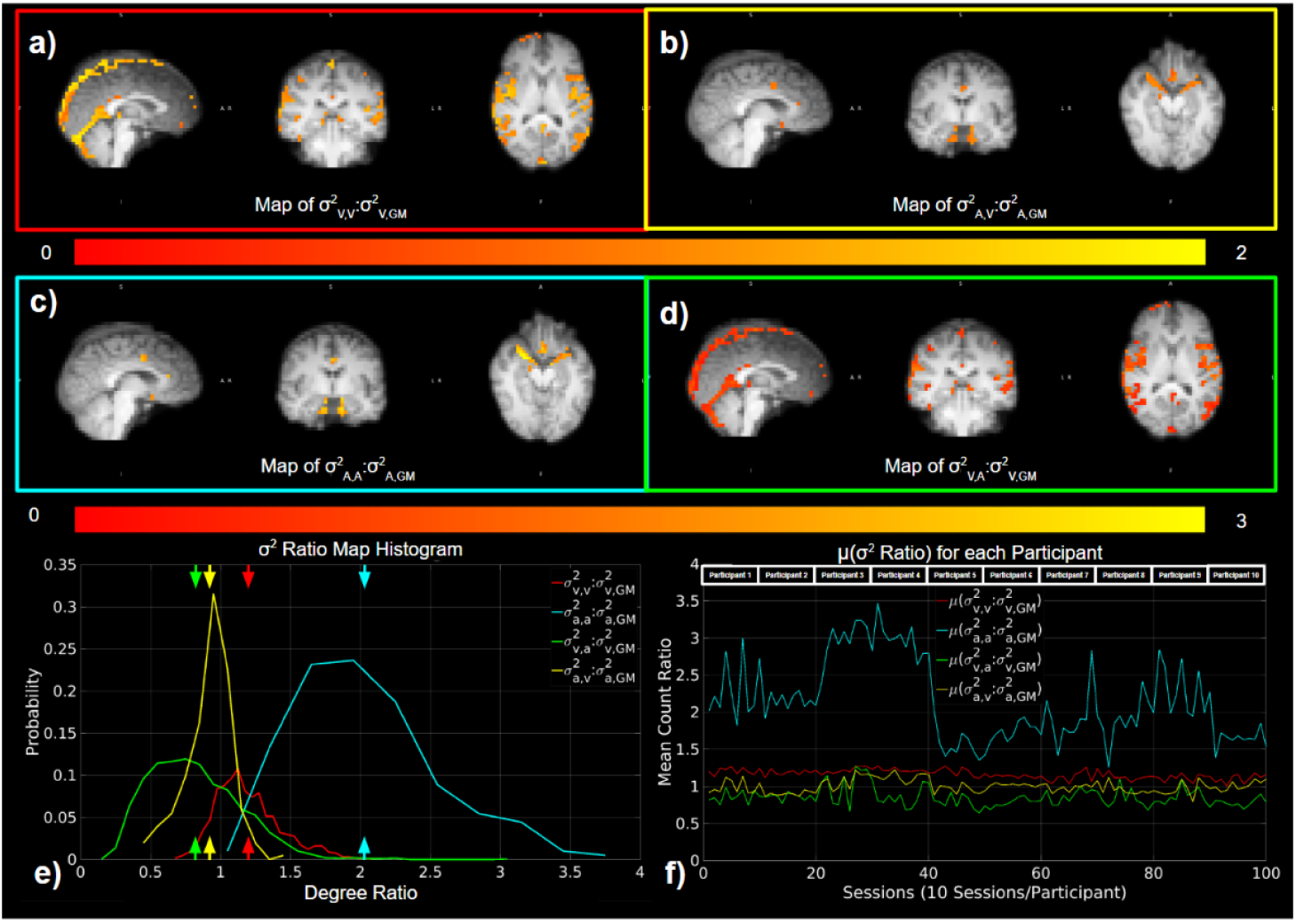
Contribution of the macrovascular connectivity to overall tissue variance of connectivity. The contribution was quantified as ratios between vascular and tissue vrainces of connectivity, e.g. σ^2^_v,v_:σ^2^_v,GM_, σ^2^_v,a_:σ^2^_v,GM_, σ^2^_a,v_:σ^2^_a,GM_, and σ^2^_a,a_:σ^2^_a,GM_. Red indicates σ^2^_v,v_:σ^2^_v,GM_; yellow indicates σ^2^_a,v_:σ^2^_a,GM_; cyan indicates σ^2^_a,a_:σ^2^_a,GM_; green indicates σ^2^_v,a_:σ^2^_v,GM_. (a)-(e) represent results from a representative data set, while (f) shows results from all data sets.

### Dependence of perivascular connectivity on the distance from vasculature

The perivascular contribution of the macrovasculature to tissue connectivity is given by degree ratios (D_A(ex),A_:D_A(ex),GM_ and D_A(ex),A_:D_A(ex),GM_) (Fig. 8) and variances ratios (σ^2^_V(ex),V_:σ^2^_V(ex),GM_ and σ^2^_A(ex),A_:σ^2^_A(ex),GM)_ (Fig. 9).

**Figure 8.**
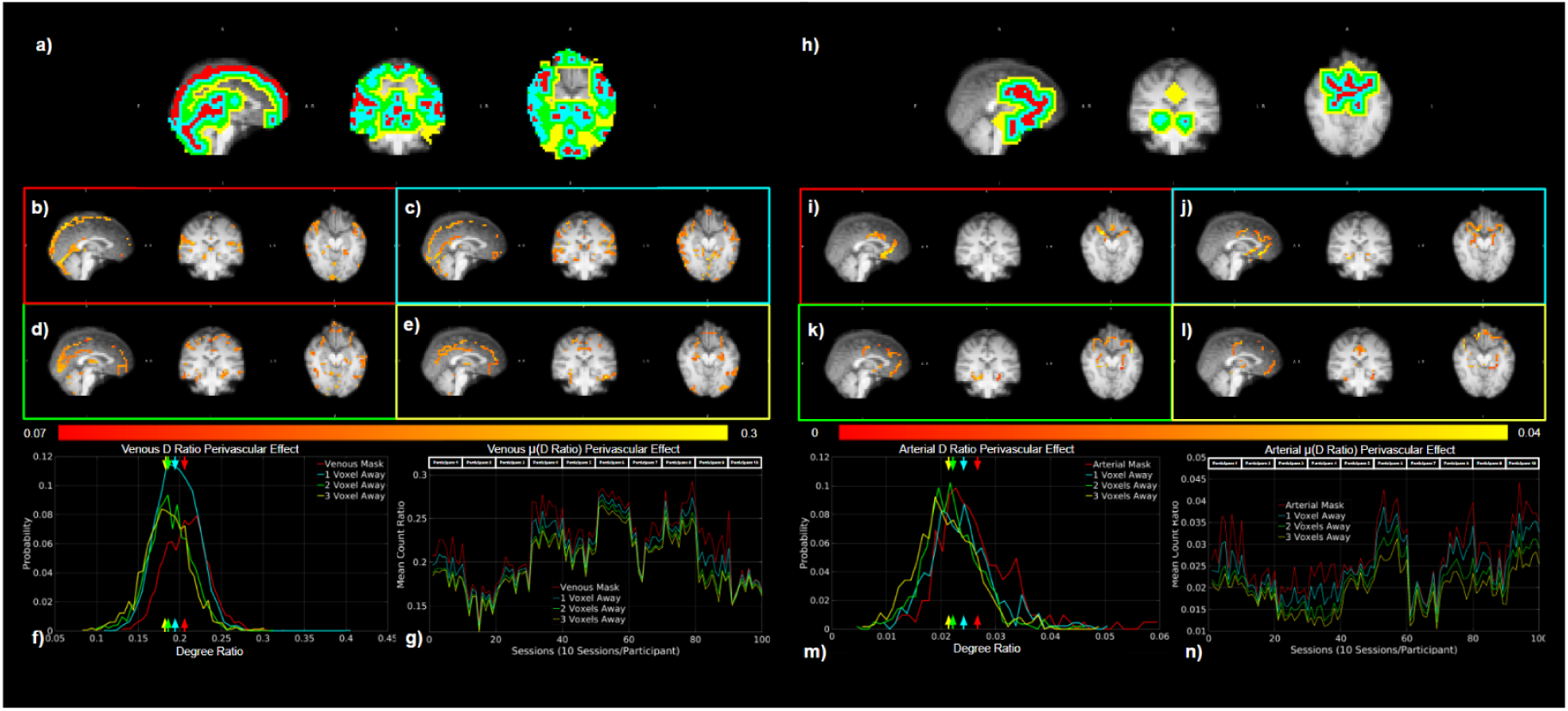
Dependence of perivascular degree of connectivity on distance from vasculature. The perivascular contribution of the macrovasculature to tissue connectivity is given by degree ratios D_a(ex),a_:D_a(ex),GM_ (a-g) and D_a(ex),a_:D_a(ex),GM_ (h-n). All plots represent results from the same representative data set, except for (g) and (n), which show results from all data sets. Extravascular ROIs with different distances to the macrovasculature are shown for both arteries and veins, with the colour coding for extravascular distance defined as: red – vasculature, blue – one voxel away, green – two voxels away, and yellow – three voxels away (a,h).

**Figure 9.**
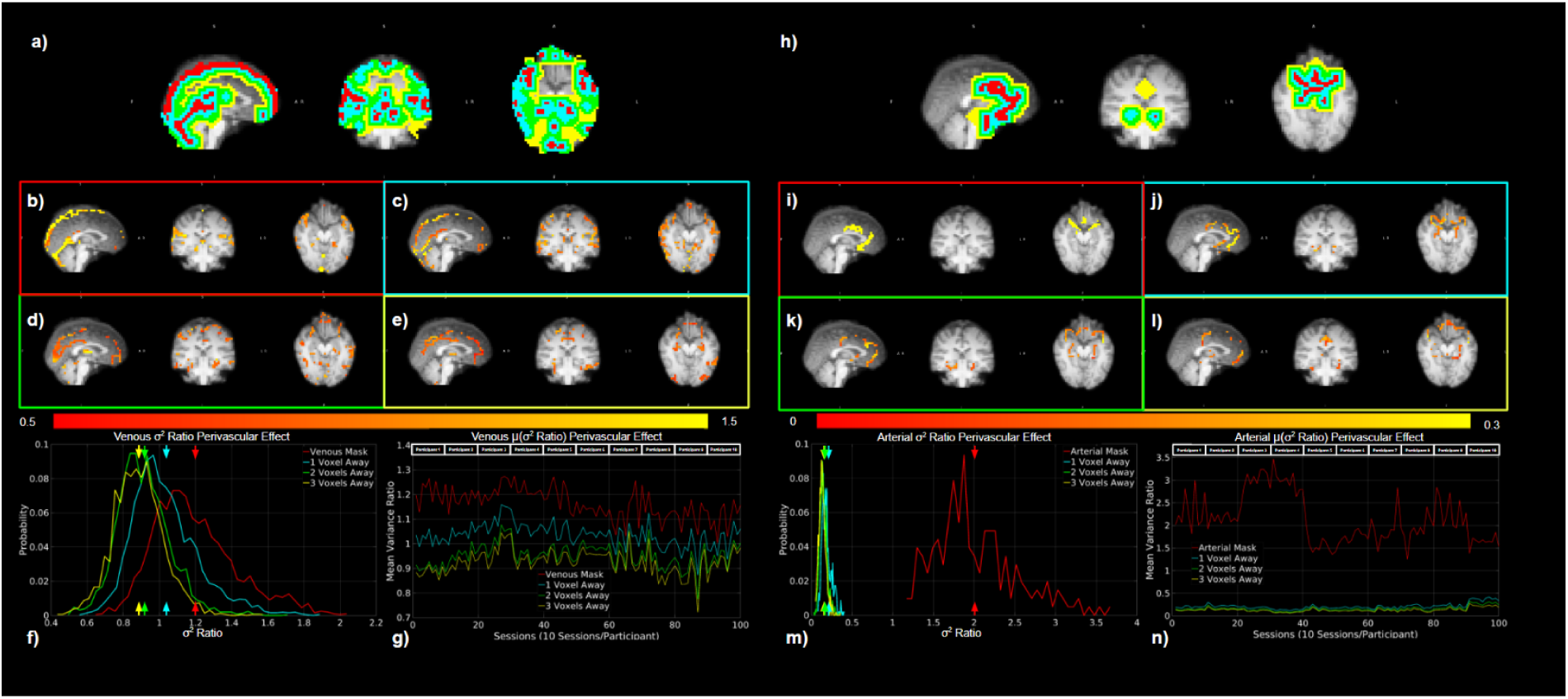
Dependence of perivascular variance of connectivity on distance from vasculature. An illustration of σ^2^_v(ex),v_:σ^2^_v(ex),GM_ (a-g) and σ^2^_a(ex),s_:σ^2^_a(ex),GM_ (h-n). Dilated vascular masks are shown for both arterial and venous, with colour coding as: red – vasculature, blue – one voxel away, green – two voxels away, and yellow – three voxels away (a,h). All plots represent results from the same representative data set, except for (g) and (n), which show results from all data sets.

### Perivascular contribution to tissue connectivity: connectivity degree

According to a representative subject as used in the previous figures, D_V(ex),V_:D_V(ex),GM_ decreases with increasing distance from the vasculature, from a maximum of 41.1% for one voxel away from the vasculature proper to a maximum of 30.6% at three voxels away (Fig. 8b**-e**). A similar trend was observed for D_A(ex),A_:D_A(ex),GM_, with a maximum of 5.11% at one voxel away, 4.69% for two voxels away, and 4.98% for three voxels away (Fig. 8i**-l**). For the perivascular degree ratios D_V(ex),V_:D_V(ex),GM_ and D_A(ex),A_:D_A(ex),GM_ across the ROIs, the histogram demonstrated the same trend and further demonstrated that the difference between the D_V(ex),V_:D_V(ex),GM_ and D_A(ex),A_:D_A(ex),GM_ of vasculature and one voxel away was higher than the difference between two voxels and three voxels away (Fig. 8f, m). As shown by the plots, the evidence above was consistent across all sessions and participants (Fig. 8g**,n**).

### Perivascular contribution of the macrovasculature to tissue connectivity: connectivity variance

Similar perivascular contribution effects were also observed on the variance ratios. Based on the results of the sample participant, both σ^2^_V(ex),V_:σ^2^_V(ex),GM_ (Fig. 9b**-e**) and σ^2^_A(ex),A_:σ^2^_A(ex),GM_ (Fig. 9i**-l**) decreased with increasing distance from the macrovascular system, with a mean of 1.04 and 0.199 at one voxel away, 0.914 and 0.147 for two voxels away, 0.883 and 0.136 for three voxels away, respectively. It is also noteworthy that μ(σ^2^_V(ex),V_:σ^2^_V(ex),GM_) was higher than μ(σ^2^_A(ex),A_:σ^2^_A(ex),GM_) except for the macrovascular ROIs (Fig. 9f**,m**). Throughout all sessions and participants, the evidence above was consistent, and the variances were negligible after one voxel away from the large arteries (Fig. 9g**,n**).

## Discussion

In rs-fMRI functional connectivity mapping, synchronized neural-related activation is assumed to be the major factor (Biswal et al., 1995; Pan et al., 2015), and there has been little investigation of functional connectivity resulting from macrovascular signal fluctuations. In our previous study, a robust correlation was observed between the internal carotid artery and the internal jugular vein, as well as arteries and arteries (Tong et al., 2019b). This work expands the previous study to include all detectable major veins as well as arteries in the brain in a systematic analysis of the macrovascular contribution to whole-GM functional connectivity. The main results are summarized in Fig. 10, demonstrating that:

1. The macrovasculature consistently exhibited strong correlational connectivity among itself, with the sign of the correlations varying between arterial and venous connectivity;
2. GM connectivity was found to have a strong macrovascular contribution, stronger from veins than arteries;
3. Functional connectivity originating from the macrovasculature displayed disproportionately high spatial variability compared to across all GM voxels;
4. Macrovascular contributions to connectivity were still evident well beyond the confines of the macrovascular space.

**Figure 10.**
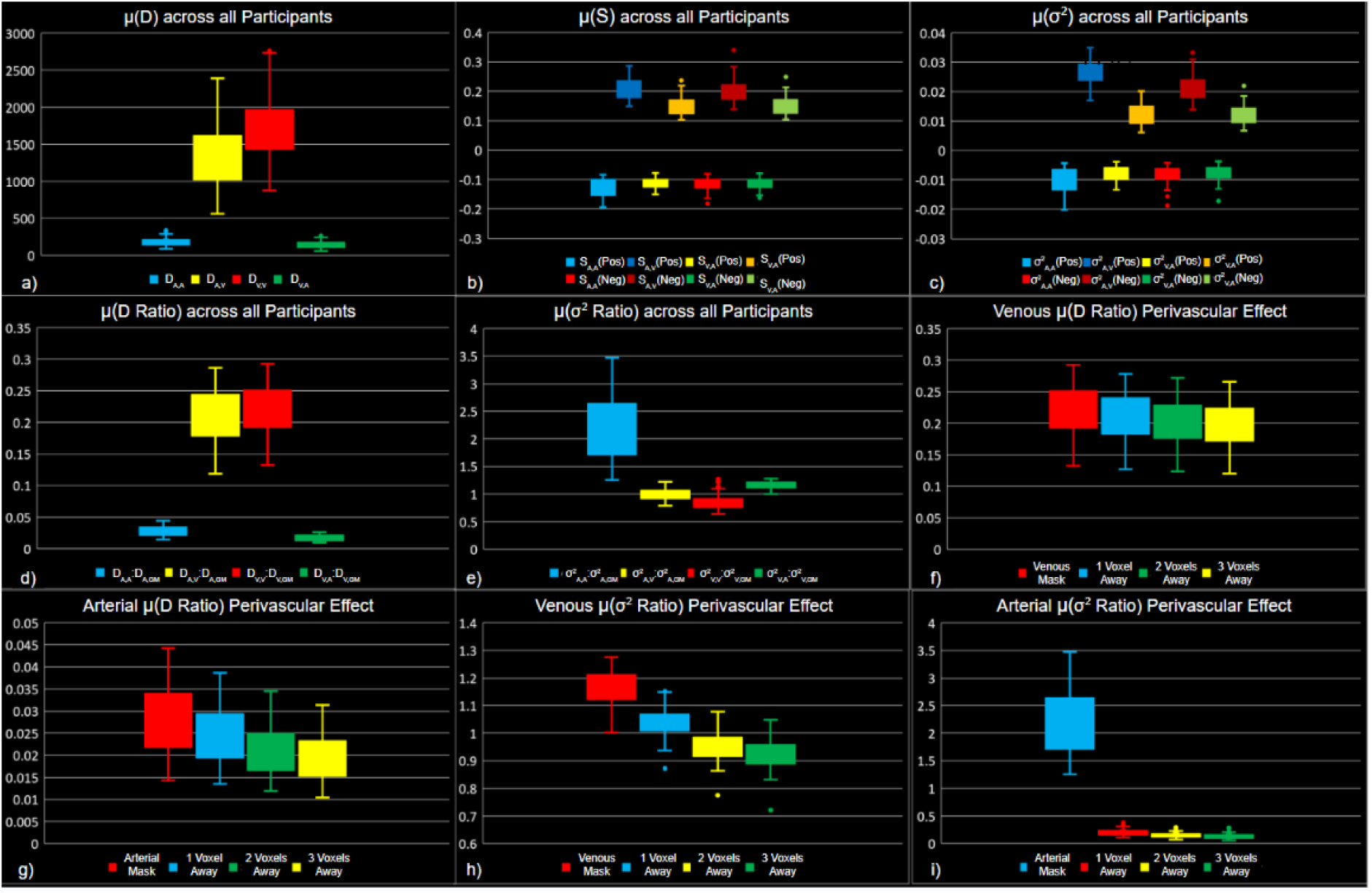
Summery box plots for each metric, with same colour encoding as Fig. 3-9. (a) Degree of connectivity; (b) mean strength of connectivity; (c) variance of connectivity; (d) ratio of degree of connectivity; (e) ratio of variance of connectivity; (f) venous perivascular ratio of degree of connectivity; (g) arterial perivascular ratio of degree of connectivity; (h) venous perivascular ratio of variance of connectivity; and (i) arterial perivascular ratio of variance of connectivity. Metrics are plotted for positive and negative correlations separately. The whiskers represent the extent of the group-wise first and fourth quartiles.

### The resting-state macrovascular BOLD effect

This is the first study to our knowledge that explores the contribution of the entire macrovasculature (to the best of our segmentation’s ability) to rs-fcMRI. In this study, we demonstrated that macrovascular contributions may bias rs-fMRI connectivity analysis, which may result in an incorrect interpretation of full brain connectivity and RSNs. An earlier review article proposed the concept of systematic low-frequency oscillations (sLFOs), which are defined as a vasogenic low-frequency BOLD signal travelling through the brain (Tong et al., 2015). It appears that the oscillation originated outside the brain (Frederick and Tong, n.d.; Li et al., 2018; Tong and Frederick, 2010) and could be caused by vasomotion (Hundley et al., 1988; Mayhew et al., 1996; Rivadulla et al., 2011), heart-rate variability (Thayer et al., 2012), respiratory volume variability (Birn et al., 2006; Chang et al., 2009), gastric oscillations (Mohamed Yacin et al., 2011; Rebollo et al., 2018) and variations in carbon dioxide levels (Sassaroli et al., 2012; Wise et al., 2004). Furthermore, previous studies using near-infrared spectroscopy have shown that such oscillations travel through the vasculature, and that strong and vascular-specific spatial structures can be captured in the macrovasculature by regressing finger-tip oxygenation time courses with the whole-brain BOLD signal (Frederick et al., 2013; Tong et al., 2015, 2014, 2011a, 2011b, 2011c; Tong and Frederick, 2014, 2012). Given the predominance of the BOLD signal by venous blood (Ogawa et al., 1993a; Segebarth et al., 1994), such sLFOs may indicate that some findings derived from rs-fMRI may be the result of venous bias rather than neural activity (Aso et al., 2020).

### Macrovascular connectivity

Our investigation of macrovascular connectivity is largely influenced by previous work by Tong et al. (Tong et al., 2015). In spite of the fact that their study only included a few very large arteries and veins, they were successful in showing a strong cross-correlation between large blood vessels, as well as between the large blood vessels and the global BOLD signal. Additionally, their results suggest that time delay control may also play an important role in macrovascular connectivity. Nevertheless, cross-correlation with different delays is not the most common method used in functional connectivity analysis, which means that whether such macrovascular effects can still be observed with simple Pearson correlation without time delays is also an important consideration. Moreover, it would be necessary to investigate whether the correlation can also be observed in the rest of the macrovasculature.

In line with the previous study discussed above, this study showed a strong correlation between BOLD fMRI signals from arterial and venous voxels (Figs. 3 and 4). As a point of clarification, our correlation results were based on Pearson’s correlation rather than cross-correlation, motivated by the stronger relevance of the former to functional connectivity mapping. Thus, unlike in Tong et al’s work (Tong et al., 2019b), arterial-venous time delays were not considered in calculating the correlation. Nonetheless, like Tong et al. we found that the correlations between venous BOLD signals were mainly positive, whereas the correlations between arterial and venous BOLD signals were mainly negative. Thus, the results demonstrated in the IJV and ICA in Tong et al.’s work are largely generalizable to other large vessels. A key difference between arteries and veins is their magnetic susceptibility, which is proportional to oxygenation and drives the amplitude of the BOLD signal (Ogawa et al., 1993b; Spees et al., 2001). At 3 Tesla, which is used in this study, the magnetic susceptibility of brain tissue and fully oxygenated blood is ∼ –9.05 ppm while that of deoxygenated blood is ∼ –7.9 ppm (Duyn and Schenck, 2017). Compared to the GM parenchyma, veins have a lower oxygenation level and contain paramagnetic blood, while large arteries have a higher oxygenation and contain diamagnetic blood. Thus we expect the intravascular BOLD-signal oscillations are naturally anti-correlated (not considering the effect of temporal delay). However, in our results, not all arterial-arterial correlations were positive, and the same was true for venous-venous correlations. These arterial-arterial and venous-venous anti-correlations were also observed by Tong et al. as shown by the histograms of the peak cross-correlation coefficients. It is possible that the phenomenon is a result of the combination of intravascular and extravascular signals. By definition, the Intervascular and extravascular signals are anti-correlated, whereas a combination of these within the same voxel could be dominated by either of these sources.

In order to provide a better global view of the macrovascular contribution to functional connectivity, we additionally computed the degree of connectivity (D) and the strength of connectivity (S) as defined in (Buckner et al., 2009; Cole et al., 2010). Calculated as the number of voxels that have high correlation coefficients above the threshold, D represents the number of voxels that may be influenced by macrovascular signals. S, on the other hand, represents the mean strength of the macrovascular contribution. In our results, we found D_v,v_ and S_v,v_ higher than D_V,A_ and S_V,A_ (or D_A,V_ and S_A,V_), which is the same as their results of superior sagittal sinus (SSS) v.s. the internal jugular vein (IJV) and internal carotid artery (ICA) v.s. IJV correlation (Tong et al., 2019b). Their correlation coefficients, however, were much higher than ours, possibly due to their use of cross-correlation and higher temporal resolution.

A notable recent study by Huck et al. found that the intravascular venous contributions to the rs-fMRI signal decreased with increasing vascular size (Huck et al., 2023). While we did not explicitly examine the effect of diameter, qualitative examination of our data do not corroborate such a finding. For instance, although the diameter of the SSS was believed to be higher than that of the straight sinus (Larson et al., 2020), the degree and strength of connectivity from the SSS were no lower than that from straight sinus (Fig. 4). However, as we did not explicitly examine the effect of vascular diameter, we cannot conclusively comment on such differences. Nonetheless, we believe that vessel diameter is only one of the required parameters to characterize the vascular contribution to the rs-fMRI signal. In theory, an analytical model that incorporates vascular diameter, oxygenation and orientation (relative to the main magnetic field), among other variables, can help reduce the inter-study and inter-subject variabilities in the apparent venous contribution. Moreover, based on the spatial distribution maps shown in Figure 3 and Figure 4, we suggest that macrovasculature connectivity might be more susceptible to changes in orientation rather than diameter. For instance, the anterior and posterior segments of SSS, for example, had weaker strength (S_V,V_ and S_V,A_) and lower degree (D_V,V_ and D_V,A_) of connectivity than the superior segment. This phenomenon might be explained by different combinations of macrovasculature-related intravascular and extravascular fluctuations as described in the biophysical model of BOLD signal (Ogawa et al., 1993b).

Beyond to these two previous studies, our study examined the connectivity within arterial vasculature, which also demonstrated a high degree and strength of connectivity (D_A,A_ and S_A,A_). In addition, we found that correlations with a high correlation strength (e.g. S_A,A_ and S_V,V_) tended to have higher correlation variance for both positive and negative correlations. This trend is also observed in spatial distributions, indicating that venous-venous correlation and arterial-arterial correlation may contribute to different RSNs to varying degrees. The macrovascular connectivity is also observable and strong at the group level, even though the extent of the effects varies among the participants.

We also noted a large amount of inter-subject variability, particularly for the degree ratio based on venous-venous correlation, arterial-venous correlation, and variance ratio based on arterial-arterial correlation. The inter-subject variability could be attributed to participants having different partial volume effects during the scan, which alter the amount of GM signal that contaminates the macrovascular signal and makes the correlation analysis less macrovascular specific. In addition, the inter-subject variability may be influenced by the brain region, which may suffer from less variability if it has a greater portion of macrovasculature. Further, the scan parameters represent potential dependencies for intersubject variability, since the temporal resolution of the imaging could affect correlation accuracy. The details of each dependency contribution would, however, be beyond the scope of this study and need future research.

### Contribution of the macrovascular connectivity to GM functional connectivity

Macrovascular connectivity is more interpretable in the context of global connectivity. The ratios of degree of connectivity, as a more direct measurement, are meant to reflect the strength of macrovasculature connectivity in relation to macrovasculature connectivity in the GM. According to our findings, macrovascular degree of connectivity constitutes a large number of significant connections when related to whole-GM degree of connectivity (a maximum of 48.8% for D_v,v_:D_v,GM_ and 7.98% for D_A,A_:D_A,GM_), even without considering the extravascular BOLD effects related to the macrovasculature. Thus, when macrovasculature is included in GM ROIs, which is often the case, up to 1/3 of the significant connections could be coming from the macrovasculature. Taken together with our previous work, which showed that connectivity z-scores decreased with increasing macrovascular blood volume fraction in a number of RSNs (Tak et al., 2015), these findings indicate the macrovasculature-driven functional connectivity is indeed of concern despite their weaker connectivity strengths compared to non-macrovascular connectivity. In addition, μ(D_V,V_:D_V,GM_) was higher than μ(D_A,A_:D_A,GM_) (Fig. 6), suggesting stronger correlations between BOLD signals from veins than from arteries (Tong et al., 2019b). This finding may also be driven by the fact veins account for a larger fraction of the GM ROI than arteries, which may also explain why μ(D_A,V_:D_A,GM_), derived from all venous voxels, is higher than μ(D_V,A_:D_V,GM_), derived from arterial voxels only.

Our findings echo findings reported by Huck et al. based on the binned values of low-frequency fluctuation amplitude and regional connectivity homogeneity, diameter and distance metrics (Huck et al., 2023). However, these results are in conflict with their voxel-based modelling results using higher-order polynomials, which fit the above GM metrics (dependent variables) against venous diameter and distance from the nearest veins (as independent variables). As the goodness of fit was used to indicate the macrovascular contribution, the latter was deemed to be lower in the presence of larger diameters. One explanation for this is that the larger variability of the effects associated with larger vessels may have diminished the ability of the polynomials to fit to the connectivity metrics. Indeed, there are other biophysical parameters that would factor strongly in producing the BOLD signal especially around larger vessels, especially vessel orientation. We have previously reported on the strong influence of vessel orientation on the BOLD signal (Zhong and Chen, 2022), so suggest that a biophysical model driven by first principles that includes the relevant variables such as vascular size, oxygenation and orientation would be needed to more fully characterize the macrovascular contribution to BOLD. Measures such as vascular position within a voxel may also strongly influence the macrovascular BOLD effect (Zhong and Chen, 2023), but are nonetheless less feasible to quantify. It is also worth noting that our results show that μ(σ^2^:σ^2^) values are much higher than the mean of other variance ratios, suggesting that the arteries have higher inhomogeneity in terms of connectivity contribution (Fig. 7). This may be due to the smaller extent of arterial vascularity compared to venous vascularity. Furthermore, the mean variance ratios of all four correlation pairs range from approximately 0.75 to 3.5, and the histograms indicate that over half of the vascular voxels exhibit connectivity variability superior to that seen in the GM. As a result of the high variance ratios, it appears that the neural specificity of GM connectivity may be heavily impacted by whether macrovascular connectivity is included in its calculation.

Interestingly, we found that increasing the threshold for degree ratio did not necessarily result in a lower degree ratio **(Fig. S1)**. For example, while we increased the threshold from 0.15 to 0.5, the results from a representative subject showed the macrovasculature still had a significant influence on the correlation between vasculature and the GM. In this case, the increase in threshold may have had less effect on the connectivity between macrovasculature and GM than on the connectivity within macrovasculature.

### Contribution of the perivascular connectivity to GM functional connectivity

Finally, we showed that the macrovascular contributions to connectivity were still evident well beyond the confines of the macrovascular space. The ratios of degree and variance were also used to determine the extent of perivascular (extravascular) connectivity. As seen in Fig. 6, the venous-to-GM degree ratio ranged between 10% and 40% even at one voxel away from the edge of the intravascular space, while arterial degree ratios were mostly less than 6%. This is in accordance with the recent results of Huck et al., who showed that major veins can produce a significant systemic spatial gradient across all common rs-fMRI metrics (including low-frequency fluctuation amplitude, regional connectivity inhomogeneity, Hurst exponent and eigenvector centrality) as a function of the distance from the veins (Huck et al., 2023). As the distance from the macrovasculature increases, the degree ratio decreases. However, the ratio of the degree of connectivity indicated that the macrovascular effect was still detectable at three voxels away from the macrovasculature, similar to reported previously (Huck et al., 2023). Considering the resolution of the rs-fMRI scans in the Midnight Scan Club dataset is 4 mm isotropic, 3 voxels amount to 12 mm. At such a low spatial resolution, macrovascular effects are likely to be present throughout the majority of GM voxels, but at a higher resolution, 12 mm may simply translate into a 6-voxel distance, still representing a large swathe of the GM. Moreover, upon dividing the macrovasculature into veins and arteries, veins (higher D_V(ex),V_:D_V(ex),GM_) exhibited greater effects than arteries (lower D_A(ex),A_:D_A(ex),GM_). As a case in point, venous connectivity can still contribute to a group-average maximum of 30.6% of whole-GM connectivity even with 3 voxels away, but arterial perivascular effects (D_A(ex),A_:D_A(ex),GM_) are as low as 6% at just one voxel away (Fig. 8f**,m**). This is in keeping with the expectation that the majority of BOLD effects are driven by the venous system (Ogawa et al., 1993a; Segebarth et al., 1994),.

Similar to the degree ratios, the connectivity variance ratios, which reflect spatial inhomogeneity of correlation strengths, decreased with increasing distance from vessels for both veins and arteries, signalling that spatial variability in functional connectivity may in large part be due to the macrovascular influence. The vascular variance ratios ranged from 0.5 to 1.5, as shown in Fig. 9, which, in conjunction with the histograms, indicate that over half of the vascular voxels exhibit connectivity variability surpassing that typically seen in the GM. Thus, if vascular or perivascular voxels are included in regional connectivity analysis, they can potentially increase connectivity variance substantially and reduce the neuronal interpretability of the connectivity measures. Similar findings pertain to the perivascular connectivity variance, although the perivascular contribution to connectivity variance diminishes with increasing distance from the vessel (Fig. 9).

It is important to note that the spatial variance of the prevalence of perivascular effect varies widely between arteries and veins. the group-average variance ratio μ(σ^2^_V(ex),V_:σ^2^_V(ex),GM_) was 1.04, 0.914 and 0.883 at one, two, and three voxels away from macrovasculature, respectively. Conversely, μ(σ^2^_A(ex),A_:σ^2^_A(ex),GM_) was only 0.199, 0.147, 0.136 for these distances, respectively (Fig. 9f**,m**). Moreover, the vascular and perivascular effect also depends on the network or region in question. For reference, we computed the mean likelihood of observing large vessels in a set of RSNs derived from the Human Connectome Project (Nozais et al., 2023), based on previously published macrovascular frequency maps (Viviani, 2016) (**Table 1**). The findings reveal that the anterior cingulate and primary visual networks contained the largest amount of macrovasculature, up to 8% even without accounting for the extravacular regions of vascular influence. Of course, the actual macrovascular contributions in these networks also depend on physical attributes of the blood vessels involved, such as vascular diameter and orientation, among others. Of note, it can be observed from Fig. 9 that the orientation of the blood vessel with respect to the B0 field may also modulate the amount of connectivity variance contributed by each blood vessel. For instance, in Fig. 9b, it is apparent that the superior-inferior segment of the sagittal sinus exhibited lower variance ratios than the anterior-posterior segment. However, the effect of vascular orientation is not explicitly modelled in this study, which focuses on the observable contributions. The biophysical modelling of such an effect will also need to take into account blood volume fraction and oxygenation, and will be the topic of our future work.

**Table 1:** List of percent macrovascular voxel occupancy well-established RSNs. This table summarizes the percentage of voxels occupied by macrovasculature for each RSN. A high percentage (above 5%) of occupancy was observed in primary visual, primary auditory, anterior cingulate, salience and cingulo-opercular areas. DMN = default-mode network. FPT = Fronto-parieto-temporal network, L = left, R = right.

| RSN Name | Percentage of RSN occupied by macrovasculature (%) |
| --- | --- |
| Anterior cingulate | 8.05 |
| Med. Post. Occ. / Primary visual | 6.93 |
| Insula / Salience | 6.55 |
| Cingulo-opercular | 5.97 |
| Primary auditory | 5.2 |
| Posterior Occipital | 4.63 |
| **Medial Occipital** | **3.13** |
| **PH-Precuneal / Parahippocampal DMN** | **3.09** |
| **Inf. Central / SM head** | **2.66** |
| **Posterior cingulate-Precuneal / Dorsal DMN** | **2.32** |
| **Precuneus / DMN proper** | **2.04** |
| **Precentral / Motor** | **1.68** |
| **Medial-frontal** | **1.52** |
| **Left Inferior frontal / Language production** | **1.42** |
| **Dorsolateral prefrontal cortex** | **1.32** |
| **Superior Parietal / Dorsal attention** | **1.23** |
| **Mid. Central (R) / SM hand (L)** | **1.00** |
| **Mid. Central (L) / SM hand (R)** | **0.89** |
| **FPT 1 (L)** | **0.87** |
| **PP-Precuneal / Posterior DMN** | **0.8** |
| **Anterior Superior Parietal** | **0.73** |
| **Superior Central / sensorimotor body** | **0.68** |
| **FPT 1 (R)** | **0.5** |
| **FPT 3 (R)** | **0.5** |
| **Posterior Superior Parietal** | **0.36** |
| **FPT 2 (L)** | **0.33** |
| **Lateral Posterior Occipital** | **0.21** |
| **Lateral Occipital** | **0.14** |
| **FPT 3 (L) / Language comprehension** | **0.14** |
| **FPT 2 (R)** | **0.13** |

Two possible explanations for the above perivascular connectivity relate to magnetic susceptibility and scan technical constraints, respectively. The magnetic susceptibility difference between the blood and tissue will generate an extravascular dipolar magnetic field offset around macrovasculature, which can extend well beyond the voxel containing the vessel (Ogawa et al., 1993b). Due to such magnetic-field offsets, it is possible for voxels in the perivascular space to experience macrovascular-like oscillations and show a high degree of connectivity with one another. On the other hand, as the BOLD signal propagates from the macrovascular system to the medium or small blood vessels, their reduced size and flow velocities may lead to challenges in capturing them in the TOF data. In spite of the fact that these vessels exhibit similar oscillation patterns to macrovasculature, demonstrate a strong correlation with macrovasculature and might have negligible time delays (if they directly connect to large vessels), they may be “invisible” to us. It is important to note that either of these factors cannot be excluded based solely on in-vivo experimental results (both degree ratios and variance ratios), especially since these two factors may interact, and future simulation studies are necessary.

### Recommendations

It is important to note that this study is focused only on assessing the macrovascular effects rather than correcting them or modelling them, which is the focus of our ongoing work. The key to removing macrovascular bias is to locate the macrovasculature. Therefore, we recommend that angiograms and venograms be collected along with rs-fMRI with either TOF imaging or susceptibility weighted imaging (SWI). At first glance, masking out the detectable macrovasculature based on angiograms may be the most straightforward method of correcting the bias. However, as we have suggested, macrovascular effect could spread well into the perivascular space, which makes direct masking inadequate as a remedy. Spin echo (SE) sequences rather than the conventional gradient-echo sequences have also been proposed that to suppress the strong macrovascular susceptibility effects. However, this option entails a sacrifice of the signal-to-noise (Menon, 2012), and the use of EPI still renders SE sensitive to large-vein effects (Ragot and Chen, 2019). More recently, BOLD image phase was also suggested as a regressor to remove macrovascular effects in task-based fMRI (Menon, 2002; Stanley et al., 2021). The method, however, assumes that all large vessels produce a phase change and that intravascular phase offsets do not contribute to the overall signal, which may not be the case in all cases (Ogawa et al., 1993b). For example, with increasing vessel size, it becomes less appropriate to disregard the intravascular phase offset and orientation effect, as discussed in our previous work (Zhong and Chen, 2023, 2022). Furthermore, Huck et al. (Huck et al., 2023) suggested using high-order polynomials to model and remove both binned intravascular and binned extravascular venous bias. However, they found their model is insufficient to remove voxel-wise venous bias in rs-fMRI. Thus, the observations in this paper prompt us to consider ways of more comprehensively modelling the vascular BOLD contributions as the first step to correcting for them.

In this context, analytical modelling (Cheng et al., 2009; Ogawa et al., 1993b) is still considered to be the best method for more accurately characterizing macrovascular effects. Based on the information provided by the angiograms and venograms (fraction of blood volume and orientation of the vessels) as well as reasonable assumptions (oxygenation level), it would be possible to generate macrovascular signals using the analytical models, and use the simulated macrovascular signal to correct macrovascular biases. The extent to which such an approach can succeed is the focus of our ongoing work.

### Study limitations

At the time of writing, the Midnight Scan Club dataset is the only public dataset that contains vascular imaging and rs-fMRI data. However, there remain limitations in the macrovascular delineation based on this data set. First, due to the limited spatial resolution of angiograms and venograms as well the large range of flow velocities across vessels of varying sizes, it is difficult to segment smaller vessels and to understand better how macrovascular factors contribute to perivascular connectivity. Secondly, the determination of arterial versus venous vessels was performed manually, allowing human error into the process, although there is no known automated alternative. Thirdly, the spatial resolution of rs-fMRI is higher than that used in most rs-fMRI studies (Raimondo et al., 2021), so it is not yet clear to what extent our results can be directly translated into a more state-of-the-art spatial resolution. Notwithstanding, the finding of large perivascular effects outside the vascular space may well translate into a spatial-distance dependent extravascular contribution, one that is in agreement with studies at higher spatial resolutions (Huck et al., 2023). Furthermore, as this study focuses on the observational perspective, the macrovascular effect is not explicitly modelled. The biophysical modelling of such an effect will also need to take into account blood volume fraction, oxygenation and vascular orientation relative to the main magnetic field, and will be the topic of our future work.

## Conclusions

To conclude, we found strong FC patterns within the macrovasculature, based on both correlations and anti-correlations (depending on the type of vasculature) at 3 Tesla. Moreover, such connectivity contributes significantly to GM FC, particularly in the vicinity of veins. Furthermore, the functional connectivity originating from the macrovasculature displayed disproportionately high spatial variability when compared to the spatial variability across all GM voxels. It important to note that, the macrovascular contribution generally extend far beyond the confines of macrovasculature, and thus cannot be removed by simple masking. A more feasible approach may involve biophysical modeling of the intra– and extravascular effects, which will be the focus of our future work.

## Acknowledgements

The authors would like to acknowledge financial support from Canadian Institutes of Health Research and the Canada Research Chairs Program (JJC).

